# Global shifts in vegetation compositional resilience over the past 8,000 years

**DOI:** 10.1101/2025.10.24.684258

**Authors:** Mengna Liao, Kai Li, Chenzhi Li, Ulrike Herzschuh, Jian Ni

**Affiliations:** College of Life Sciences, Zhejiang Normal University, Jinhua 321004, China; State Key Laboratory of Lake and Watershed Science for Water Security, Nanjing Institute of Geography and Limnology, Chinese Academy of Sciences, Nanjing 211135, China; Polar Terrestrial Environmental Systems, Alfred Wegener Institute Helmholtz Centre for Polar and Marine Research, Potsdam 14473, Germany

**Author notes:** Corresponding authors. (ML), (KL).

**Keywords:** pollen analysis, early warning signals, vegetation resilience, anthropogenic land-use, climate change, diversity, synchrony, multi-millennial timescale

## Abstract

Vegetation resilience is critical for sustaining biodiversity and ecosystem services, yet its long-term dynamics and underlying drivers remain poorly resolved. Here, we assess changes in continental-scale vegetation compositional resilience over the past 8,000 years by applying established leading indicators (including autocorrelation, standard deviation, skewness, and kurtosis) to 482 fossil pollen records from all major landmasses except Antarctica. Using a machine learning approach, we evaluate the relative contributions of anthropogenic land cover change (ALCC) and climatic parameters (annual mean temperature, temperature seasonality, annual precipitation, and precipitation seasonality) in driving resilience shifts across continents. We further employ structural equation modeling (SEM) to disentangle how abiotic factors (ALCC and climatic variables) and biotic factors (species richness, evenness, synchrony, and temporal *β* diversity) jointly shape long-term trends in vegetation resilience at the continental scale. Our results reveal a persistent, millennia-scale decline in resilience during the past 1.6 to 4.4 thousand years before present, with intensified anthropogenic land-use change identified as the dominant global driver. This finding suggests that the global decline in vegetation resilience observed over recent decades likely represents a continuation of the long-term trend established during the preceding Holocene. Notably, North America exhibited a resurgence in resilience over the past 1,200 years, primarily attributed to enhanced resilience in tundra and savanna ecosystems. SEM analyses uncover complex pathways through which abiotic and biotic predictors interact to shape millennial-scale resilience changes across continents. Despite the complexity of these underlying mechanisms, the analyses demonstrate that biotic factors collectively exert substantial, and often predominant, influences on long-term resilience dynamics. This underscores the indispensable role of biotic attributes in modulating long-term vegetation resilience, highlighting their potential as critical targets for ecosystem conservation and restoration.

## Introduction

Ongoing global changes have profoundly impacted Earth’s ecosystems, forcing abruptly and irreversibly regime shifts in some ecosystems (Cavanaugh et al., 2019; Smol et al., 2005; Wernberg et al., 2016). At the planetary scale, evidence suggests that the biosphere as a whole may be approaching a catastrophic bifurcation point (a ‘tipping point’) due to sustained loss of resilience (Barnosky et al., 2012). Given that current rates of change in key Earth System features already match or exceed those of abrupt geophysical events in the past (Steffen et al., 2018), assessing the global-scale resilience of Earth’s ecosystems has become an urgent priority for developing adaptive management strategies.

Resilience was defined as the capacity of a system to resist disturbance, recover to its original state after a disturbance, and sustain fundamental functions at pre-disturbance levels (Folke et al., 2004). It represents two stability paradigms, namely local stability and non-local stability (Dakos & Kéfi, 2022). The local stability, also termed engineering resilience, typically refers to an ecosystem’s capacity to return to its original state following a small disturbance when near a stable equilibrium state (Pimm, 1984). The non-local stability, widely known as ecological resilience, can evaluate the magnitude of disturbance that can be absorbed before the system changes state (Holling, 1973). Fundamentally, engineering resilience focuses on preserving the recovery rate of a structure and/or function, while ecological resilience emphasizes maintaining the existence of the structure and/or function (Holling, 1996). Over the past two decades, ecologists proposed generic early warning signals (or leading indicators) to bridging the gap between the intuitive conceptual clarity of ecosystem resilience and its operational, quantifiable metrics (Carpenter & Brock, 2006; Dakos & Kéfi, 2022; Guttal & Jayaprakash, 2008; Scheffer et al., 2009; Wang et al., 2012). Resilience reduction, which is often associated with phenomena recognized in multi-stable ecosystems as “critical slowing down” or “flicking” can be quantitatively characterized by increases in leading indicators such as variance, autocorrelation, skewness, and kurtosis of system state variables (Dakos et al., 2008; Dakos et al., 2013; Scheffer et al., 2009; Wang et al., 2012). These phenomena can theoretically be assessed within any temporally and spatially bounded system (Barnosky et al., 2012). As different individual metric may have different predictive ability, the combination of several leading indicators into a single composite metric could enable more reliable inference of resilience changes (Drake & Griffen, 2010).

Vegetation unquestionably forms the most crucial component of terrestrial ecosystems, which provides essential ecosystem services while also maintaining the balance of the Earth System (Duveiller et al., 2018). Recent studies, employing either individual leading indicators or composite resilience indices, have demonstrated a widespread decline in vegetation resilience across many regions over the past decades (Boulton et al., 2022; Feng et al., 2021; Forzieri et al., 2022; Smith et al., 2022), based on satellite-derived vegetation metrics such as Vegetation Optical Depth (VOD) and Normalized Difference Vegetation Index (NDVI). While these metrics serve as indicators of gross primary production (Liu et al., 2015) and vegetation greenness or canopy coverage (Myneni et al., 1997), they poorly reflect vegetation composition, which is a critical dimension for understanding shifts in ecosystem structure and function disturbed by climate change and human activities. Moreover, satellite-derived vegetation metrics are limited to a temporal coverage of only several decades, which is exceedingly brief when compared to the extensive history of human-induced landscape alterations and the long-term process of vegetation succession. In addition, vegetation’s response to climate change manifests across a range of timescales from hundreds to tens of thousands of years, potentially showing non-linear characteristics or demonstrating temporal lags (Fastovich et al., 2025). Therefore, the limitations in data observation post significant challenges for empirical data in detecting critical shifts of vegetation composition resilience and attributing these shifts.

Fossil pollen records offer long-term perspectives for inferring past plant community properties and their dynamics across local, regional, and global scales spanning the last few centuries to millions of years (Birks, 2019; Jackson & Blois, 2015). By compiling pollen data, recent studies have revealed that vegetation composition has been transformed globally by climate changes and increasing human disturbances since the last glacial maximum (Gordon et al., 2024; Nolan et al., 2018). Moreover, the rates of vegetation compositional change accelerated unprecedently during the late Holocene in both magnitude and extent (Mottl et al., 2021). Given the profound transformations of vegetation ecosystems caused by climate changes and, particularly, human disturbances, large-scale vegetation resilience could have undergone significantly shifts during the Holocene. However, as of now, there is no evidence to support this hypothesis, nor has it been confirmed whether human disturbances play the crucial role in driving continental or global changes in vegetation compositional resilience.

Ecosystems not only make direct responses to external forcings but also possess inherent ecological feedbacks that enable them to mediate responses to exogenous stress, thereby sustaining the resilience of ecosystems (Blois et al., 2013; Folke et al., 2004). The stabilizing effect of biodiversity (generally measured by species richness) has received the most extensive attention and discussion (Ives & Carpenter, 2007; Loreau & De Mazancourt, 2013). Most biodiversity–stability relationships involve some forms of compensatory dynamics or insurance effect, which require either asynchrony of species responses to environmental fluctuation or the presence of functionally redundant species with differing sensitivities to specific perturbations (de Bello et al., 2021; Loreau & De Mazancourt, 2013; Oliver et al., 2015). While this certainly appears to be the case at the community level in some ecosystems over relatively short ecological timescales spanning years to no longer than decades (Hatton et al., 2024; Hisano et al., 2018; Isbell et al., 2015), these patterns may not generalize to larger spatial scales or longer temporal scales (Peterson et al., 1998; Wang & Loreau, 2014). Other biological attributes, such as species evenness and spatial self-organization, regulate ecosystem resilience as well (Kéfi et al., 2024; Valencia et al., 2020). While, the applicability of these stability mechanisms to explaining long-term resilience trajectories in vegetation composition remains uncertain. This knowledge gap underscores an urgent need to extend current knowledge to broader spatial extents and longer temporal baselines, thereby aligning research scales with the operational dimensions of global biodiversity management and conservation strategies.

Here, we assessed continental-scale vegetation resilience over the past 8,000 years using fossil pollen sequences from the recently published global late Quaternary pollen dataset (Li et al., 2022). A total of 482 pollen records (Figure 1) were selected from all continents, with the exception of Antarctica. We calculated a composite resilience index (CRI) for each record by integrating four leading indicators (*i*.*e*., autocorrelation, standard deviation, skewness, and kurtosis). The estimated CRI sequences were aggregated by continent, and each continental CRI dataset was fitted using generalized additive models (GAMs) to reconstruct long-term trajectories of vegetation compositional resilience. We then evaluated the relative importance of abiotic (including climate variables and land-use change) and biotic factors (including diversity and synchrony) to vegetation resilience, as well as their direct and indirect effects, employing a machine learning approach and piecewise structural equation model (SEM). With this study, we aim to provide a long-term perspective on trends in vegetation resilience across continental to global scales and provide new insight into the mechanisms underlying long-term vegetation resilience dynamics.

**Figure 1.**
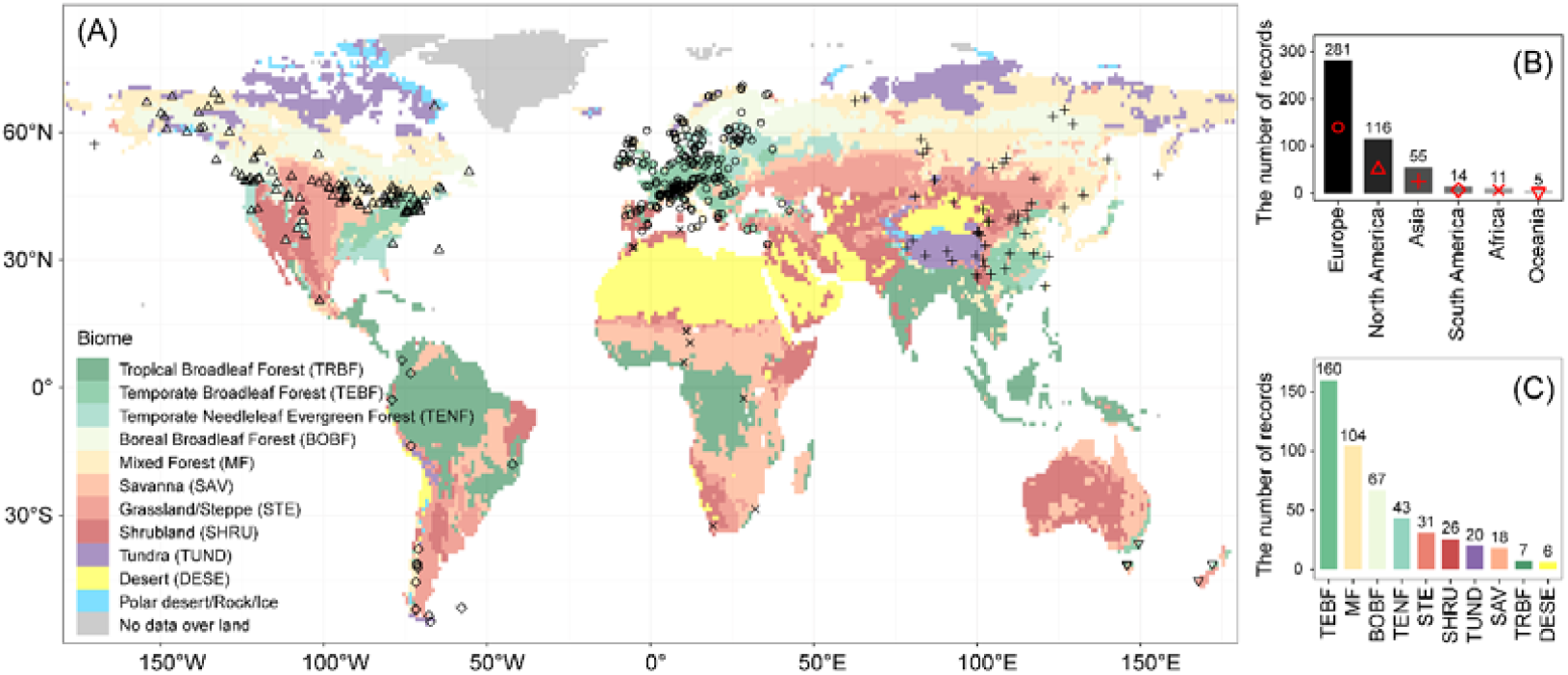
Spatial distribution of the fossil pollen sequences analyzed in this study, with sequence counts by continent and biome. (A) Spatial distribution of pollen sequences. (B) The number of pollen sequences per continent [symbols match those in (A)]. (C) The number of pollen sequences per biome [colors match those in (A)].

## Methods

### Fossil pollen data source

Fossil pollen data were obtained from the LegacyPollen 2.0 dataset, a globally harmonized compilation of 3680 palynological records (Li et al., 2024, 2025). To ensure analytical consistency across diverse records, the dataset underwent rigorous taxonomic standardization—woody and dominant herbaceous taxa were harmonized at the genus level, while other herbaceous taxa were standardized at the family level (Herzschuh et al., 2022). Temporal standardization was achieved by constructing robust Bayesian age–depth models (Blaauw & Christen, 2011; Li et al., 2022). Each record contains both percentage and count data for identified taxa, along with calibrated age estimates corresponding to sediment depth.

### Data filtering

We applied several criteria for data filtering, including archive type, the youngest and oldest depositional ages of records, the mean temporal resolution of samples, the total numbers of chronological control points, the presence or absence of sedimentary hiatus, and the temporal gap between adjacent samples. Firstly, we excluded records collected from marine, ice, and animal midden, as these archives rarely allow clear defining of the geographical source of fossil pollen. Regarding the depositional ages, we retained records with a minimal age no older than 2 cal ka BP and a basal age no younger than 10 cal ka BP. The adoption of this criterion ensures that the timeseries of the calculated vegetation resilience (see below for how CRI was calculated) spans at least from the mid-to the late Holocene. We used a mean temporal resolution of 200 years as another data filtering criterion, based on a comprehensive assessment of both data availability and the capacity of pollen data to well resolve long-term vegetation dynamics over multi-millennial timescales. To minimize uncertainties arising from potential bias in estimated depositional ages, we excluded records containing fewer than three independent chronological control points from the analysis. The presence of pronounced sedimentary hiatus or large chronological gaps between adjacent samples in pollen records may result in either misinterpretation or failure to reconstruct vegetation dynamics accurately. Given this, we calculated the chronological gaps between each pair of adjacent samples in all records that met the aforementioned criteria. If a record contains a chronological gap exceeding 2,000 years, only its upper section is retained for subsequent analysis, contingent upon this section meeting the aforementioned criteria. Ultimately, we identified 482 pollen sequences that met all the filtering criteria, including a total of 56,939 pollen samples. Among these sequences, 60.5% contain samples that predate the onset of the Holocene (11.7 cal ka BP), 19.7% extend back to 15 cal ka BP, and only 3.1% reach as far back as 20 cal ka BP (Figure S1).

### Compositional resilience index (CRI)

CRI can be computed as the sum of the standardized differences of each employed resilience metric (Drake & Griffen, 2010). In this study, we applied four leading indicators of ecological resilience namely, autocorrelation at-lag-1 (AR1), standard deviation (SD), skewness (SK), and kurtosis (KURT), due to their positive performance (Dakos et al., 2024) to compute CRI for each selected pollen record (Eqs. 1–9) (Dakos et al., 2024; Dakos et al., 2012; Drake & Griffen, 2010).

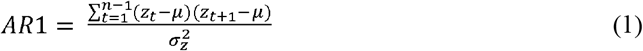

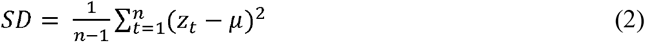

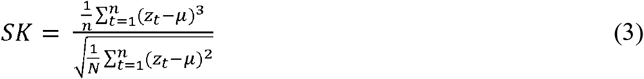

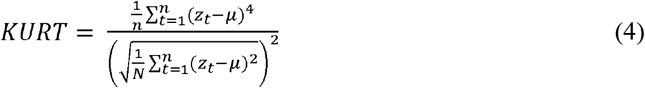

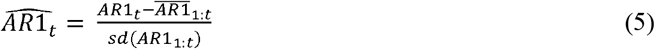

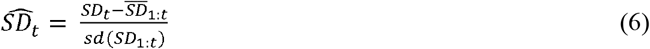

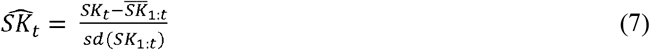

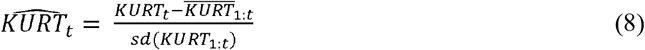

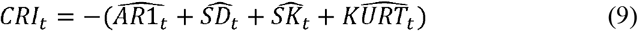

where *z*_*t*_ is the subset of time-detrended ecological state series (*i*.*e*., scores of DCA1 in this study) within a sliding window; *z*_*t+1*_ is the first-order lag series; *μ* and σ_*z*_^2^ are the mean and variance of *z*_*t*_, respectively. *n* is the size of the sliding window. *AR1*_*t*_, *SD*_*t*_, *SK*_*t*_, *KURT*_*t*_, and *CRI*_*t*_ are the *AR1, SD, SK, KURT*, and *CRI* at time *t*, respectively. *AR1*_*1:t*_, *SD*_*1:t*_, *SK*_*1:t*_, and *KURT*_*1:t*_ denote the timeseries of *AR1, SD, SK*, and *KURT* from times 1 to *t*, respectively. 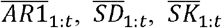, and 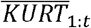 are the mean values of *AR1, SD, SK, KURT*, and *CRI* from times 1 to *t*, respectively. 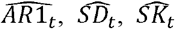, and 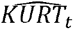 indicate the normalized *AR1, SD, SK*, and *KURT* at time *t*. Given that an increase in these leading indicators theoretically signifies a decline in system resilience, a rising in the composite index (calculated as the sum of standardized differences across all employed leading indicator) would similarly indicate reduced resilience (Drake & Griffen, 2010). To enable the composite index more intuitively reflect the direction of resilience change, we defined the *CRI*_*t*_ as the negative value of the summed standardized differences. Specifically, an increasing CRI trend indicates enhanced vegetation resilience, whereas a decreasing CRI trend signifies declined vegetation resilience.

Given that these leading indicators are applicable exclusively to one-dimensional timeseries (Dakos & Kéfi, 2022), we performed detrended correspondence analysis (DCA) using ‘decorana’ function from the ‘vegan’ package (Oksanen et al., 2022) to generate an aggregate measure reflecting the compositional state of multi-dimensional pollen assemblages. The first axis of DCA (hereafter DCA1), which generally captures the dominant gradient of compositional variation, was used as the aggregate measure for calculating the leading indicators. Additionally, an underlying assumption for applying these leading indicators is that the observations or timeseries must be equidistant. Therefore, we linearly interpolated the DCA1 scores at 200-year intervals (corresponding to the maximum mean temporal resolution of the samples) to generate equivalent timeseries (Dakos et al., 2008). Subsequently, we detrended the interpolated DCA1 scores using a locally weighted regression smoother (LOESS) to remove long-term trends. The residuals between interpolated and fitted values of DCA scores were then used to compute the leading indicators (Wang et al., 2012).

The choice of the rolling window size is a trade-off between the time-resolution of the available data and reliability of the estimate for the indicators (Lenton et al., 2012). To determine the optimal window size, we estimated AR1, SD, SK, and KURT using rolling window sizes ranging from 10% to 80% of the timeseries length, incremented in steps of five data points. Trends in the leading indicators across different window sizes were quantified using the Kendall tau rank correlation coefficient. For each indicator, the optimal rolling window size was independently selected for individual DCA1 residual timeseries based on the maximum Kendall tau value. Given that the optimal rolling window sizes generally vary across the four leading indicators for each residual timeseries, the temporal coverage of the composite index is determined by the common overlapping period among all four indicators. Eventually, continental CRI sequences for North America, Europe, and Asia span the past 8,000 years, whereas those for South America, Africa, and Oceania cover the past 7,200 or 7,000 years (Figure S1).

Leading indicator calculations and trend estimations of statistical moments with different rolling window sizes were performed using functions ‘genetic_ews’ and ‘sensitively_ews’, respectively, in ‘earlywarnings’ package (Dakos et al., 2012).

### Richness and evenness

Richness was quantified as the effective number of taxa (*Hill* number of order *q* = 0) in each pollen assemblage, while evenness was measured by assessing the equality in the abundances of individual pollen types within the assemblage (Birks et al., 2016; Hill, 1973). Given that empirical estimates of *Hill* numbers tend to be an increasing function of sampling effort or sample completeness (Chao et al., 2014), richness was determined by calculating the expected number of taxa in samples rarefied to a fixed size of 300 pollen grains. Evenness was estimated by using the modified *Hill*’s ratio (Eq. 10) (Alatalo, 1981):

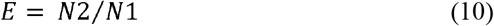

where E is evenness; *N1* is the *Hill* number of order *q* = 1 (equivalent to the exponential of Shannon entropy or diversity measure), serving as an estimate of the number of abundant taxa in an assemblage; *N2* is the *Hill* numbers of order *q* = 2 (equivalent to the inverse of Simpson’s measure index of concentration), providing an estimate of the number of very abundant taxa in an assemblage. *Hill* numbers (*q* = 0, 1, 2) were calculated using ‘iNEXT’ function from the ‘iNEXT’ package (Chao et al., 2014; Hsieh et al., 2022).

### Temporal *β* diversity

Temporal *β* diversity between temporally adjacent sample pairs within each pollen record was quantified using the Sørensen pair-wase dissimilarity, calculated via the ‘beta.pair’ function in the ‘betapart’ package (Baselga et al., 2018). To guarantee the accuracy of *β* diversity estimations along a temporal gradient, it is essential to maintain a uniform time interval between adjacent samples. For this, pollen data from each record were linearly interpolated to generate a dataset with evenly spaced interval prior to calculating temporal *β* diversity. Dataset interpolation was performed using the function ‘interp.dataset’ from the package ‘rioja’ (Juggins, 2022), with an interpolated interval set to 200 years to match the temporal resolution used for computing the CRI. Given that the algorithm for calculating the Sørensen pair-wase dissimilarity is exclusively applicable to present/absent data, all non-zero values in the interpolated pollen dataset were converted to 1.

### Synchrony

We defined synchrony between two species or taxa as the extent of correlation in their abundances over a specified time period, with values ranging from +1 (perfect synchrony) to 0 (maximum asynchrony). For a multispecies community, community-wide synchrony can commonly be measured as the mean correlation between the abundance of each species versus the rest of the community (Gross et al., 2014) or the arithmetic mean correlation coefficient across all species pairs (Loreau & de Mazancourt, 2008; Thibaut & Connolly, 2013). Given that pollen data are presented as either the percentage abundance of each taxon in samples or the grain count of each taxon within rarefied samples with equal size, it is inappropriate to apply Gross’s synchrony metric (Gross et al., 2014), as the correlation between the percentage abundance or grain count of each taxon and the rest of the assemblage would exhibit perfect correlation. Therefore, we considered community-wide synchrony metric based on correlation coefficient across all species pairs to be a feasible approach for reflecting the overall synchrony in pollen assemblages. In this study, we developed a new measure of overall synchrony for pollen assemblages as

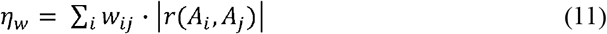

where *η*_*w*_ denotes the weighted synchrony index; *A*_*i*_ and *A*_*j*_ represent the timeseries of percentage abundance for taxon *i* and taxon *j* (*i*≠*j*), respectively; *r* is the correlation coefficient between *A*_*i*_ and *A*_*j*_ with a significant level < 0.05; *w*_*ij*_ is the sum of the mean percentage abundances of taxa *i* and *j* within an assemblage over the targeted period, namely the sum of the averages of *A*_*i*_ and *A*_*j*_. We used Kendall’s tau coefficient to quantified the correlations between paired taxa, and the significance of these correlations was evaluated using a two-tailed *t*-test. To obtain temporal trajectory of synchrony change, *η*_*w*_ was calculated within a five-point rolling window (1000 years) of the interpolated pollen dataset (200-year resolution).

### Resilience trends at continental and biome scales

We modeled continental and biome CRIs against time with GAMs. GAMs are nonparametric data-driven regression models, which are effective in assessing non-linear relationships between response and predictor variables without any restrictive assumptions (Hastie & Tibshirani, 1987). To facilitate model convergence, we standardized each CRI timeseries using z-score normalization, which involves subtracting the mean from each data point and then dividing by the standard deviation. The standardized CRI timeseries were subsequently aggregated into continental subsets and biome subsets according to their geological locations. The information on the continents where the pollen records were obtained is from the LegacyPollen2.0 dataset. The ISLSCP II Potential Natural Vegetation dataset, sourced from the Earthdata platform (https://www.earthdata.nasa.gov/) and comprising fifteen vegetation types, was applied as the reference for defining biome subsets. For each subset, we fitted a GAM to the pooled standardized CRI values (hereafter, CRI_st) using the ‘gam’ function in the ‘mgcv’ package (Wood, 2011). In this process, we estimated a smooth function representing the relationship between CRI_st and time, assuming a Gaussian distribution. When formulating GAM equation, we designated CRI_st as the response variable and used the age corresponding to each CRI_st value as the predictor variable. We incorporated a thin plate regression spline smoother into the formula, setting the smoothing basis dimension to five for representing the smooth term. Given the heterogeneity (*e*.*g*., variations in environmental conditions, local ecological differences) across sampling sites from which pollen records were derived, we included a random effect of sampling sites into the model to provide a more realistic representation of the underlying data structure and yield more robust results. The significance of the GAMs was evaluated using the *F* statistic.

### Investigating key drivers for continental vegetating resilience shifts

We used a random forest (RF) model to identify the primary driver(s) responsible for the transition of continental resilience from an increasing to a declining trend during the late Holocene. RF analysis is a flexible, non-parametric regression tool which can handle a large number of predictors even in small datasets with a low number of observations, and is suited to analyzing non-linear relationships and complex interactions (Feld et al., 2016). To capture the states of resilience trends, we created a dummy variable based on the fitted continental-scale trends of CRI_st prior to conducting the analysis. Specifically, we assigned a value of 1 to periods showing an increasing trend and 0 to those showing a decreasing trend. It is worth noting that, given our focus on the increase-to-decline transition during the late Holocene, we excluded data from the dummy variable for two specific periods: (1) post-1.4 cal ka BP in North America, where trajectories deviated from the post-transition decline, and (2) pre-5.8 cal ka BP in South America, where trajectories deviated from the pre-transition increase (Figure 2). This approach allowed us to eliminate potential bias in assessing the relative importance of predictors that may arise from the existence of other transitions in the CRI_st trends. Additionally, RF analysis was not applied to Africa due to the absence of a discernible transition in its CRI_st trend (Figure 2).

**Figure 2.**
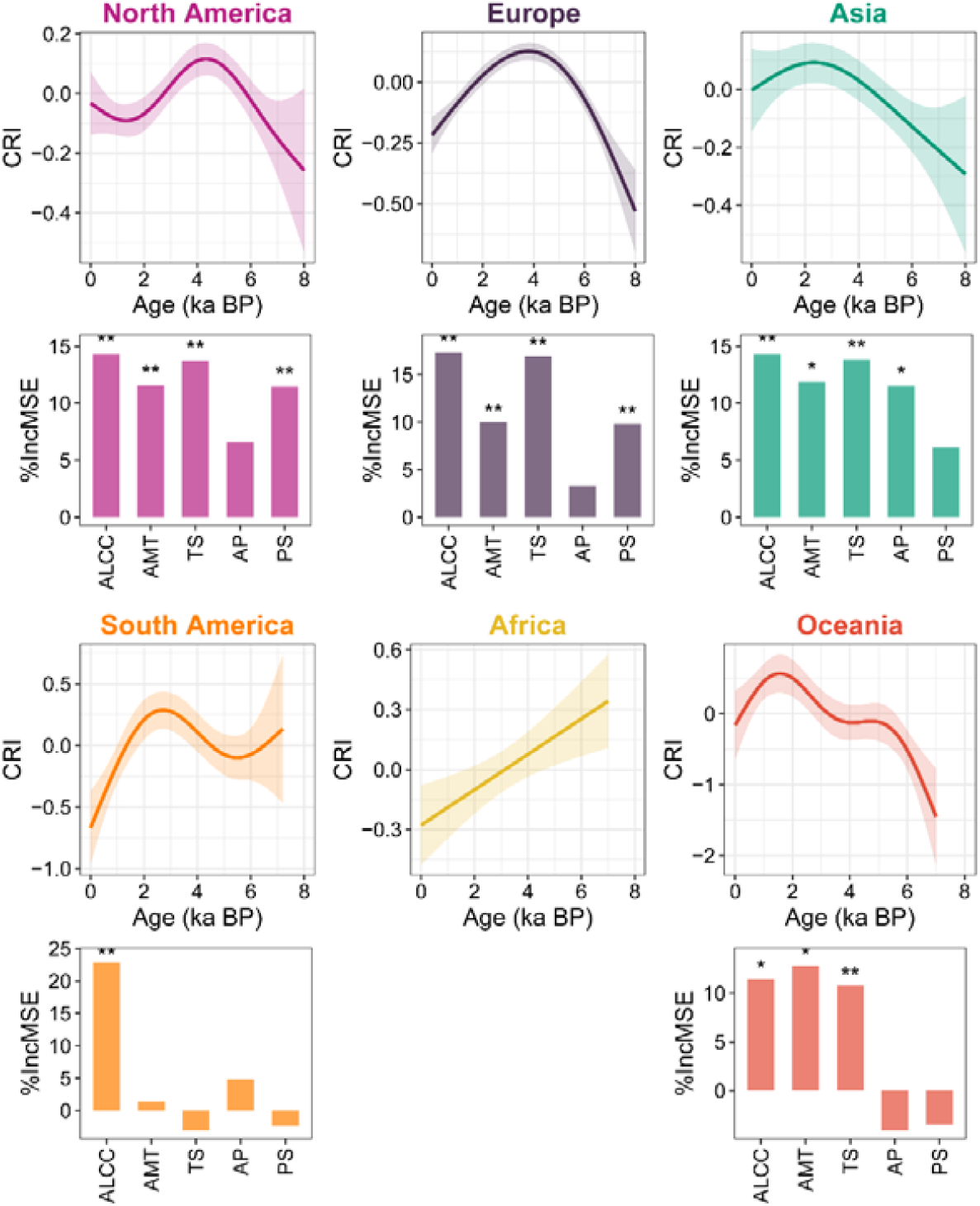
Continental CRI trends and relative importance of abiotic drivers. Solid lines denote significant fits (*P* < 0.05) of generalized additive model (GAM), while dashed lines indicate non-significant relationship (*P* > 0.05). The shaded areas represent the 95% confidential intervals of the GAM fits. Histograms show the relative importance of each driver, quantified as percentage increase in the mean squared error (%*InMSE*), with significant drivers marked with asterisks (*P* < 0.05 and ** *P* < 0.01). ALCC, anthropogenic land-cover change; AMT, annual mean temperature; TS, temperature seasonality; AP, annual precipitation; PS, precipitation seasonality.SEMs showing the direct and indirect effects

In RF models, we used the dummy variable representing the resilience state as the response variable. The predictor variables included annual mean precipitation (AMT), temperature seasonality (TS), annual precipitation (AP), and precipitation seasonality (PS), which were derived from CHELSA-TraCE21k dataset (Karger et al, 2023), as well as anthropogenic land use, which was obtained from the KK10 dataset scenario of Anthropogenic Land Cover Change (ALCC) (Kaplan et al., 2011). Based on the geographical coordinates of the pollen records, we extracted timeseries data of the predictor variables for all pollen record from the two datasets using the function ‘extract’ from the package ‘terra’ (Hijmans, 2023). The time slices for the extracted data were selected to match those of the CRI_st timeseries. To generate continental-scale timeseries for each predictor variable, we firstly standardized each predictor’s timeseries using z-score transformation and then partitioned the standardized timeseries into six continental subsets based on their geographical locations (Figure S2). For each predictor, continental-scale timeseries were generated by computing the median values across all time slices within each continental subset.

Variable importance was quantified by the increase in out-of-bag error when the values of the variable are randomly permuted (Breiman, 2001). We estimated the importance of predictor variables using the percentage increase in the mean squared error (%*InMSE*), where higher %*InMSE* values indicate greater variable importance (Breiman, 2001). The significance of each predictor was assessed with 1,000 permutations of the response variable. A predictor was considered statistically significant if the permutation-induced increase in *MSE* exceeded the 95^th^ percentile of the null distribution generated from these permutations. RF analyses and the permutation were conducted using the functions ‘rfPermute’ and ‘importance’ from the package ‘rfPermute’ (Archer, 2023).

### Attributing long-term changes of continental resilience

We performed piecewise structural equation models (SEMs) based on a maximum likelihood method to test direct and indirect pathways of biotic and abiotic factors on continental-scale resilience of vegetation composition. SEM provides a way to test postulated relationships between causal and confounding variables and to disentangle their effects (Shipley, 2016). We constructed an initial SEM including climatic variables (AMT, TS, AP, PS) and ALCC as observed variables, biotic attributes (richness, evenness, temporal *β* diversity, synchrony) as latent variables, and the CRI as the dependent variable. The initial structure contained all possible biotic and abiotic relationships with CRI, but did not define structural paths among latent variables. Standardized path coefficients were used to measure the direct and indirect effects of predictors (Grace & Bollen, 2005). We refined the model structure by adding potentially significant pathway(s) and/or removing those with extremely weak standardized path coefficient to improve the fit. The minimal and best model was selected based on the Akaike Information Criterion (AIC). Fisher’s C test was conducted to assess the global goodness-of-fit of the models (*P* ≥ 0.05 indicated reliable models). We conducted SEM analyses using the function ‘psem’ from the package ‘piecewiseSEM’ (Lefcheck, 2016), performing separately for each continent to examine continental-specific causal relationships among variables. Following a similar data processing protocol to that used for RF analysis, we generated continental-scale timeseries for each variable through multi-step procedure involving standardization of data using z-score normalization, partitioning of standardized data into continental subsets, and calculating median values for each time slices within each subset (Figure S3).

In SEM, direct effects were measured as the standardized path coefficients between variables, while indirect effects were calculated as the product of direct path coefficients along a specific causal chain. We calculated the direct and indirect effects for each observed and laten variable, independent of the statistical significance of the path coefficient(s). The net effect of each variable was estimated by summing its direct and indirect effects. In addition, we classified the observed and laten variables into three types: climate, ALCC, and ecological attributes, to evaluate the relative contributions of different factor types on resilience. The net effects of each factor type were quantified as the sum of all direct and indirect effects of the variables grouped within that type. Then, we evaluated the relative effect sizes of distinct factor types for each continental-scale SEM. Effect size was defined as the absolute value of the net effect. The relative contributions of each factor type were quantified as the proportion (%) of its effect size relative to the total effect size across all factor types.

We performed all data manipulations and analyses using R (Team, 2022). All packages mentioned in this study are software extensions to the R environment.

## Results

### Continental-scale variations in vegetation resilience

All continents except Africa show significantly reversed CRI trends over the past 8,000 years (Figure 2). Notably, these reversed trends predominantly comprise a pattern of increase during the Mid-Holocene [8.0 to 4.2 thousand calibrated years before radiocarbon present (cal ka BP, where 0 ka BP is 1950 CE)], followed by a decline in the Late Holocene (4.2 to 0 cal ka BP), although the onset of these declines varied across continents (Figure 2). The earliest decline signal was detected in North America (4.4 cal ka BP), followed by Europe where the CRI began to decrease at 3.6 cal ka BP. For South America, Asia, and Oceania, the declines in CRI trends occurred at 2.8, 2.4, and 1.6 cal ka BP, respectively. Interestingly, in North America, the CRI trajectory over the past 1,200 years deviates from the “increase–decrease” pattern, showing an increasing trend instead.

The main abiotic predictors driving the transition of continental-scale CRI trends from increasing to declining during the late Holocene were identified through RF analysis (Figure 2). ALCC was the most important variable for predicting the transition across all continents, except Oceania where AMT surpassed ALCC as the top-ranked driver. In addition to ALCC, the trend transition of CRIs in North America and Europe was significantly influenced by TS, PS, and AMT, whereas in Asia it was significantly affected by TS, AMT, and AP. In Oceania, TS was another significant predictor that had considerable impact on the CRI shift.

### Pathways and effects of abiotic and biotic attributes on continental-scale vegetation resilience

Pathways and effects of abiotic factors (AM, TS, AP, PS, ALCC) and ecological attributes (taxonomic richness, evenness, temporal *β* diversity, and synchrony) on CRIs have been estimated by SEMs (Figure 3). In North America, the most prominent path is the direct link from ALCC to CRI. Evenness plays a significant role in influencing CRI, serving both as a significant direct driver to CRI variations and as a mediator that relays the effects of other factors to CRI. TS and richness have significant impacts on evenness, subsequently further affecting CRI. Moreover, ALCC exerts an indirect influence on CRI through richness. In Europe, temporal *β* diversity is the only factor that displays a significant direct path influencing the CRI. Richness shows a significant indirect path to CRI through temporal *β* diversity. ALCC exerts a substantial influence on richness, which subsequently further impacts CRI. In Asia, TS displays a strong positive direct path to CRI, while PS shows a strong negative direct path to CRI. Furthermore, synchrony exhibits a significant negative direct link to CRI, through which ALCC exerts negative effects on CRI. Similar to Europe, there are no significant direct paths from biotic factors to CRI, and temporal *β* diversity is the only factor that shows a significant direct path influencing CRI in South America. In this SEM, abiotic factors contribute to CRI via several significant indirect paths, including those from AMT and evenness via temporal *β* diversity, from ALCC, PS, and richness via evenness, and from ALCC via richness. The SEM for Africa reveals only one significant direct path originating from AP. Similar to Africa, the significant direct paths originate solely from abiotic factors in Oceania; however, the abiotic factors exerting significant impacts are ALCC and AMT, rather than AP.

**Figure 3.**
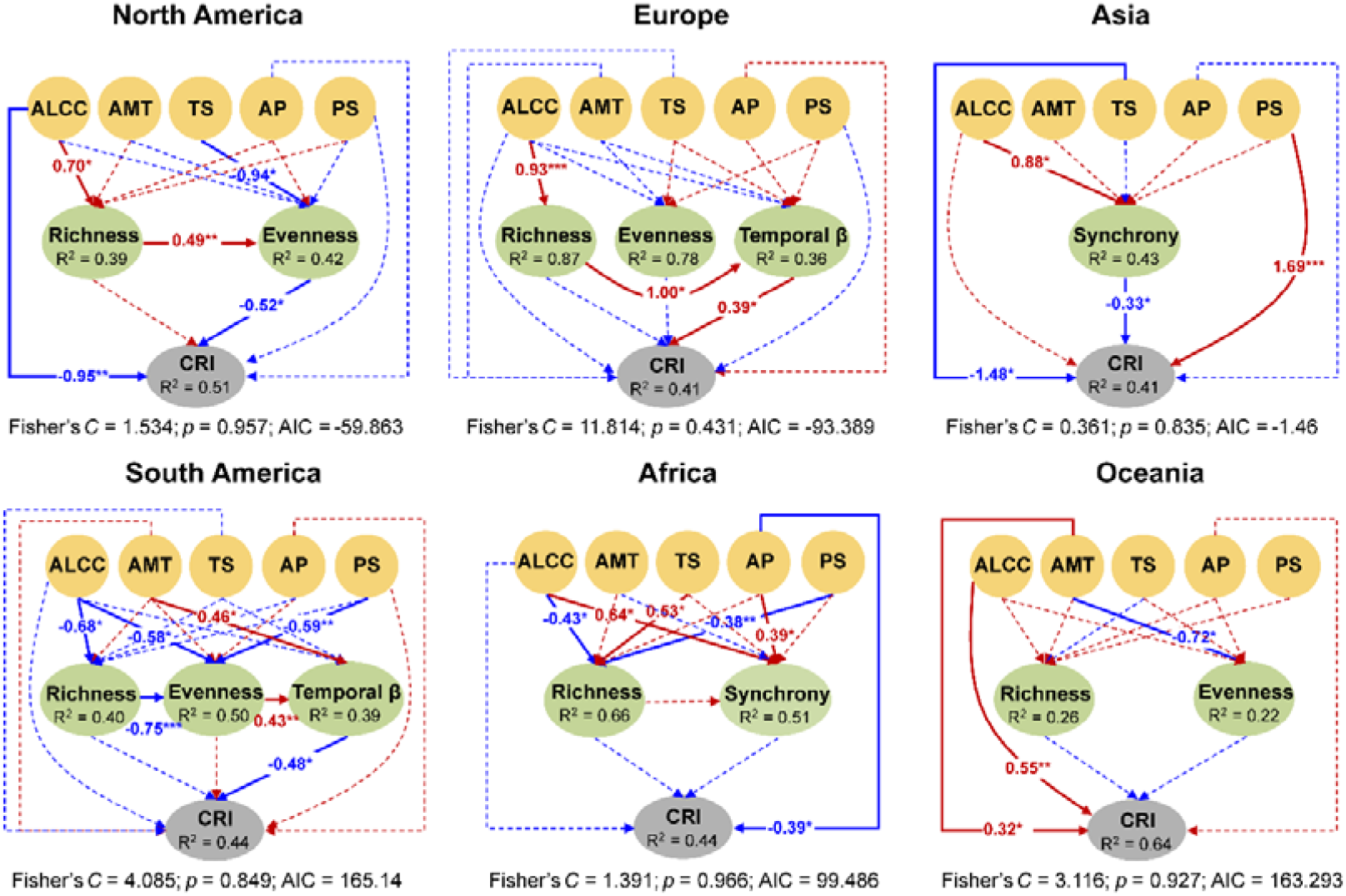
SEMs showing the direct and indirect effects of multiple abiotic and biotic drivers on CRI across six continents. Solid lines denote statistically significant paths (*P* < 0.05), while dashed lines indicate non-significant paths. Red and blue lines represent positive and negative effects, respectively. *R*^2^ values indicate model-explained variance for each response variable. Standardized path coefficients are reported only for significant paths with significant level marked (* *P* < 0.05, ** *P* < 0.01, *** *P* < 0.001). Overall fit of SEM was evaluated using Shipley’s test of d-separation: Fisher’s *C* statistic (if *P* > 0.05, no paths are missing and the model is a good fit) and Akaike information criterion (AIC). ALCC, anthropogenic land-cover change; AMT, annual mean temperature; TS, temperature seasonality; AP, annual precipitation; PS, precipitation seasonality.

The factors incorporated into the SEMs show varying effect sizes across different continents (Figure 4). In North America, evenness contributes most to CRI variations, with net effect size (*n*.*e*.*s*., thereafter) of 0.7057. Although richness demonstrates both substantial positive and indirect negative effects on CRI, its net effect is weak. ALCC exerts a relatively strong effect on CRI in North America (*n*.*e*.*s*. = 0.1579), while other abiotic factors contribute comparatively minimal influence to CRI variation. In Europe, temporal *β* diversity (*n*.*e*.*s*. = 0.4274) emerges as the strongest driver of CRI variation, followed by evenness (*n*.*e*.*s*. = 0.3028). In Europe, the *n*.*e*.*s*. of richness on CRI is weak and ALCC is identified as the strongest abiotic driver of long-term CRI variation (*n*.*e*.*s*. = −.2745). In Asia, PS (*n*.*e*.*s*. = 0.5886) and TS (*n*.*e*.*s*. = 0.54) show the notably stronger effects on CRI variation than all other evaluated factors. Synchrony also contributes a substantial effect on CRI (*n*.*e*.*s*. = −0.511). In South America, richness (*n*.*e*.*s*. = −0.5452) and temporal *β* diversity (*n*.*e*.*s* = −0.5142) are two strongest predictors of CRI variations, and AP (*n*.*e*.*s*. = 0.2169), ALCC (*n*.*e*.*s*. = 0.2023) and PS (*n*.*e*.*s*. = 0.1973) are the most important abiotic drivers. In Africa, richness (*n*.*e*.*s*. = 0.4255) and synchrony (*n*.*e*.*s*. = 0.3274) show obvious larger effect sizes on CRI variations than all other evaluated factors. Among abiotic factors, ALCC (*n*.*e*.*s*. = 0.1781) is the strongest predictor of CRI variation in Africa, following by AP (*n*.*e*.*s*. = −0.1563) and TS (*n*.*e*.*s*. = 0.1811). AMT (*n*.*e*.*s*. = 0.4119) and ALCC (*n*.*e*.*s*. = 0.3216) are the two most important factors influencing CRI in Oceania. Among biotic factors, the effect of evenness (*n*.*e*.*s*. = 0.3692) is stronger than that of richness (*n*.*e*.*s*. = 0.2069).

**Figure 4.**
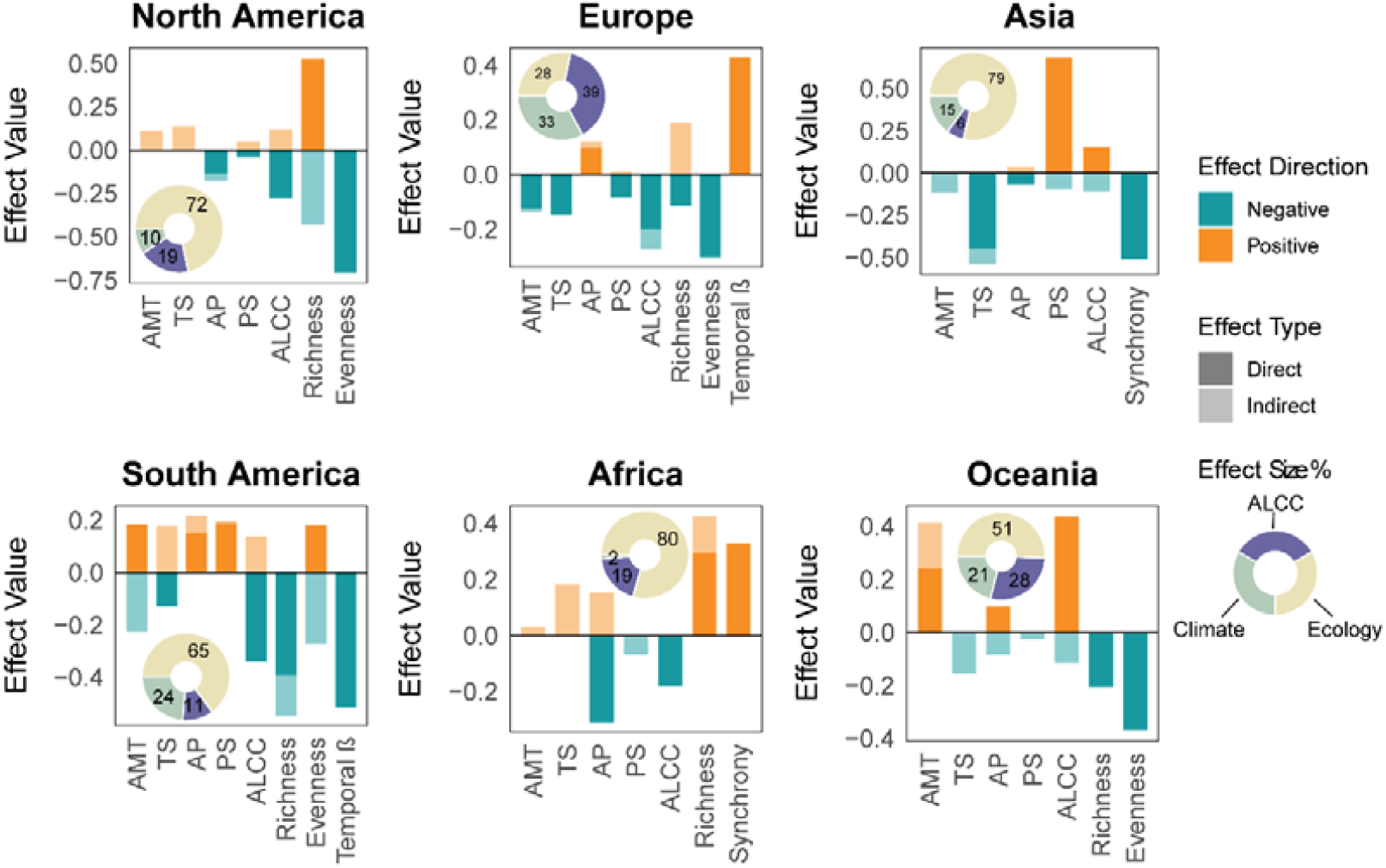
Effect sizes of abiotic and biotic drivers on continental CRI trajectories. Donut charts illustrate the relative net effect of each grouped variable to continental CRI changes. ALCC, anthropogenic land-cover change; AMT, annual mean temperature; TS, temperature seasonality; AP, annual precipitation; PS, precipitation seasonality.

When quantifying the total net effects, ecological attributes account for a predominant proportion of the total *n*.*e*.*s*., ranging from *ca* 51% to *ca* 80%, across all continents except Europe (Figure 4). In Europe, the three contributors of ALCC, climate (*i*.*e*., AMT, TS, AP, and PS), and ecological exhibit relatively comparable *n*.*e*.*s*., with ALCC emerging as the most important factor. Although evenness and temporal *β* diversity have the largest *n*.*e*.*s*. among all evaluated factors in the SEM for Europe, their opposing effects diminish the overall impact of ecological attributes on CRI. A similar situation emerged in Asia, where PS and TS are the two most influential variables affecting CRI, yet their counteracting effects result in climate contributing only a minor portion of the total net effect on CRI. When comparing abiotic factors, climate exerts a notable stronger influence than ALCC on CRI variations in South America and Asia. Conversely, ALCC plays a more important role than climate in North America and Africa, while the *n*.*e*.*s*. of these two contributors are comparable in Europe and Oceania.

## Discussion

Pollen archives have recorded the global shifts of vegetation compositional resilience during the Holocene, with the onsets of resilience declines occurred approximately between 4.4 and 1.6 cal ka BP across different continents except Africa (Figure 2). This aligns with that the Holocene Earth experienced significant vegetation changes, manifesting in multiple dimensions such as shifts in the compositional and structural in terrestrial vegetation (Nolan et al., 2018), accelerated rates of compositional change (Mottl et al., 2021), and alterations in floristic diversity (Gordon et al., 2024). However, the continental-scale trends in resilience differ from those observed in other ecosystem properties, including composition (Figure S4), turnover rates (Mottl et al., 2021), and biodiversity (Gordon et al., 2024). In addition, transitions in these properties often occur asynchronously across continents and/or biomes, reflecting the inherent non-linearity and heterogeneity of ecosystem responses to Holocene environmental changes. Interestingly, the global resilience decline reversed in North America at *ca* 1.2 cal ka BP (Figure 2). This phenomenon can be collectively attributed to the shifts in the resilience trajectories of steppe, tundra, and savanna biomes during the past 1,500 and 2,000 years, as well as the concurrent slowdown in the resilience loss in forest and shrubland in North America (Figure S5). On a global scale, compositional resilience in tundra and savanna ecosystems diverged from their preceding declining trajectories and initiated a recovery at around 3–2 cal ka BP (Figure S6). Thus, the continental-to-global scale shifts in vegetation resilience should be predominantly shaped by the integrated effects of different ecosystem responses to external stress, with such differential responses likely contingent upon the intrinsic properties of these ecosystems (Pennekamp et al., 2018).

ALCC was primary factor that triggered the reversed trends of continental vegetation resilience during the late Holocene (Figure 2). This was particularly evident in South America where ALCC emerged as the sole statistically significant predictor driving reverse in the CRI trend. Although human modification on Earth’s landscapes can be traced back to the Paleolithic era, its adverse effects on vegetation compositional resilience did not occur until the late Holocene. Among the diverse array of resource procurement and land management, intensive agriculture’s high demand for resources and its continuous modification of land makes it the strongest force in changing the Earth’s landscapes. Intensive agriculture was prevalent only in a few regions with suitable climatic conditions until 4–3 cal ka BP, after which it began to spread more widely across the globe (Stephens et al., 2019). The rapid expansion of intensive agriculture has largely coincided with the decline in the compositional resilience of global vegetation, suggesting the existence of a land-use threshold inducing shifts in the long-term trajectories of continental vegetation resilience. In addition to ALCC, climatic variables contributed substantially to the shifts in the long-term trends of vegetation resilience (Figure 2), suggesting that the observed resilience decline was commonly trigged by the synergistic effects of human disturbance and climate change.

Both theoretical and empirical researches have delved into how various features of ecosystems, such as complexity of ecosystems (Pimm, 1984), biodiversity (Liang et al., 2025; Pennekamp et al., 2018; Rodrigues et al., 2025; S. Wang & Loreau, 2016), species synchrony (Schnabel et al., 2021; Valencia et al., 2020), species evenness (Isbell et al., 2009; Valencia et al., 2020), spatial self-organization (de Paoli et al., 2017; Kéfi et al., 2024), affect ecosystem stability/resilience. The relationship between biodiversity and stability has long been a contentious issue, primarily focused on the question of whether greater diversity (number of species) leads to less variability in communities. Our findings reveals that species richness can exert positive, negative, or negligible effects on vegetation resilience, at least in terms of vegetation composition, when assessed across continental scale and millennial timescales (Figure 3). This conclusion supports that a general conclusion about the diversity-stability cannot be anticipated (Ives & Carpenter, 2007), since external pressures often affect stability and diversity simultaneously. The stabilizing effects of biodiversity arise from species’ asynchronous responses to environmental fluctuations, a phenomenon commonly referred as insurance effects (Loreau et al., 2021; Yachi & Loreau, 1999). However, species synchrony was identified as an important structural component of the SEMs only for Asia and Africa (Figure 3). In Asia, the SEM demonstrates a negative link between synchrony and vegetation compositional resilience, which appears to align with the insurance effect. In contrast, the SEM of Africa reveals a positive pathway between synchrony and resilience. Furthermore, we observe a positive richness-synchrony relationship in Africa, which appears to contradict the prevailing hypothesis that higher species richness enhances the likelihood of a community containing species with differing responses to abiotic drivers or competition, thereby reducing synchrony (McCann, 2000). However, the positive correlation between species richness and synchrony observed in this study aligns with findings from an empirical global assessment that examined how species richness and synchrony influence community stability (Valencia et al., 2020). These findings collectively suggest that the effect of richness on stability/resilience is not necessarily mediated through its negative influence on synchrony, either at the local scale or at the continental scale.

Our study identifies evenness as another important direct or indirect predictor of long-term changes in vegetation compositional resilience across North and South America, Europe, and Oceania (Figure 3). Theoretically, declines in evenness may decrease temporal stability/resilience by weakening the portfolio effect or statistical averaging (Doak et al., 1998; Hillebrand et al., 2008). While, the SEMs predominantly reveal a negative association between evenness and resilience (Figure 3). This is likely because, analogous to the mass-ratio hypothesis (Grime, 1998), dominant species exert stronger effect on vegetation dynamics than common and rare taxa do. When a vegetation ecosystem operates under a background where the dominant taxa progressively lose their ecological dominance (*i*.*e*., evenness increase), a gradual decline of composition stability or resilience will arise from the reduced involvement of the dominants in shaping ecosystem structure and dynamics. Temporal *β* diversity, defined as temporal variation in community composition or species assemblages at a specific location (Magurran et al., 2019), comprises two distinct components: species turnover (resulting from species replacement) and nestedness (resulting from species loss or gain) (Baselga, 2010). In this study, temporal *β* diversity is quantified using adjacent samples from time series with fixed temporal intervals, thereby serving as a metric for the rate of compositional change in an assemblage. It was expected that the variations in compositional change rates were expected to influence vegetation resilience, and this hypothesis is verified by the SEMs for Europe and South America (Figures 3 & 4). However, similar to species richness and evenness, the direction of temporal *β* diversity’s impact on resilience is not fixed.

Despite these complexities present above, there are some general underlying mechanisms that regulate changes in vegetation compositional resilience. Our study reveals that the assessed abiotic factors exert certain impacts on the biotic factors, and that some or all of these abiotic factors have direct influences on resilience (Figure 3). These findings emphasize that biotic factors unlikely change in isolation from abiotic drivers influencing stability, and in fact these abiotic drivers are typically the main causes of changes in ecological attributes and may serve as the original driving force for variations in ecosystem stability or resilience (Ives & Carpenter, 2007). As another generality, both biotic and abiotic factors may have opposing effects on resilience through their direct and direct pathways (Figure 4). In some cases, a certain factor may appear to have a negligible impact on resilience because it simultaneously generates positive and negative effects of comparable magnitude. However, this does not necessarily imply that the positive and negative effects produced by this factor are weak *per se*. Regarding the relative importance of different factor types, biotic factors as a whole generally exert a substantially greater influence on continental-scale changes in compositional resilience than abiotic factors do, although Europe represents an exception to this pattern (Figure 4). Furthermore, while ALCC has been identified as the primary driver for the shift in the long-term trend of vegetation compositional resilience during the late Holocene, its cumulative effect on resilience trajectories over the past 8,000 years has not exceeded the cumulative influence of climate change across all continents.

In conclusion, our results suggest that human-induced modifications to Earth’s landscapes have led to a persistent, global-scale decline in vegetation resilience that has lasted for several millennia. However, we have also found that over the last 1,000–2,000 years, in some regions constrained by moisture and/or temperature conditions, such as tundra and savanna ecosystems, vegetation resilience has continuously enhanced. Regarding the mechanisms driving millennia-scale changes in continental vegetation resilience, this study reveals a complex picture when incorporating factors representing human disturbance, climate changes, and ecological attributes. Despite this, the results demonstrate that biological attributes generally contribute more than other factors in regulating the long-term changes in vegetation resilience. There are two limitations in our data that could potentially affect the findings presented here. First, the sparse pollen records in Africa and Australia may lead to a lack of precision when analyzing the long-term trends of vegetation resilience in these two continents. In addition, since pollen data generally lack high taxonomic resolution, in cases where the changes in vegetation composition primarily manifest as alternations in plant taxa within the same genus or family, the variations in vegetation composition are likely to be underestimated. This may subsequently bias the estimated resilience metrics based on taxonomic compositional changes, thereby affecting accurate assessments of resilience dynamics.

## Acknowledgements

We acknowledge the contributions of all data providers who supported this study. We appreciate Professor Rong Wang (Nanjing Institute of Geography & Limnology, Chinese Academy of Sciences) for his insightful discussions on methodological approaches. This work was supported by the National Natural Science Foundation of China (No. 42477484, No. 42577519) and the State Key Laboratory for Vegetation Structure, Function and Construction (No. VegLabOF2025006).

## Author Contributions

**Mengna Liao:** Conceptualization (lead); data curation (equal); formal analysis (lead); funding acquisition (equal); methodology (lead); software (lead); supervision (equal); validation (lead); visualization (lead); writing – original draft (lead); writing – review and editing (equal). **Kai Li:** Conceptualization (supporting); funding acquisition (equal); writing – original draft (supporting); writing – review and editing (equal). **Chenzhi Li:** Data Curation (equal); investigation (equal); writing – review and editing (equal). **Ulrike Herzschuh:** Data Curation (equal); investigation (equal); writing – review and editing (equal). **Jian Ni:** Conceptualization (supporting); investigation (equal); supervision (equal); writing – review and editing (equal).

## Data Availability and Code Availability

The LegacyPollen2.0 pollen dataset is openly available at https://doi.pangaea.de/10.1594/PANGAEA.965907. The CHELSA-TraCE21k climate data is openly available at https://www.chelsa-climate.org/datasets/chelsa-trace21k-centennial. The KK10 Anthropogenic Land Cover data is openly available at https://doi.pangaea.de/10.1594/PANGAEA.871369. The R scripts are available on GitHub at https://github.com/mnliao/Global-shifts-vegetation-resilience.git.

**Supplementary Materials** was provided as a separated PDF file.

