## Supplemental Figures S1 to S6 for "Global shifts in vegetation compositional resilience over the past 8,000 years": Supplementary Materials_updated.pdf

**This PDF file includes:** Figures S1 to S6

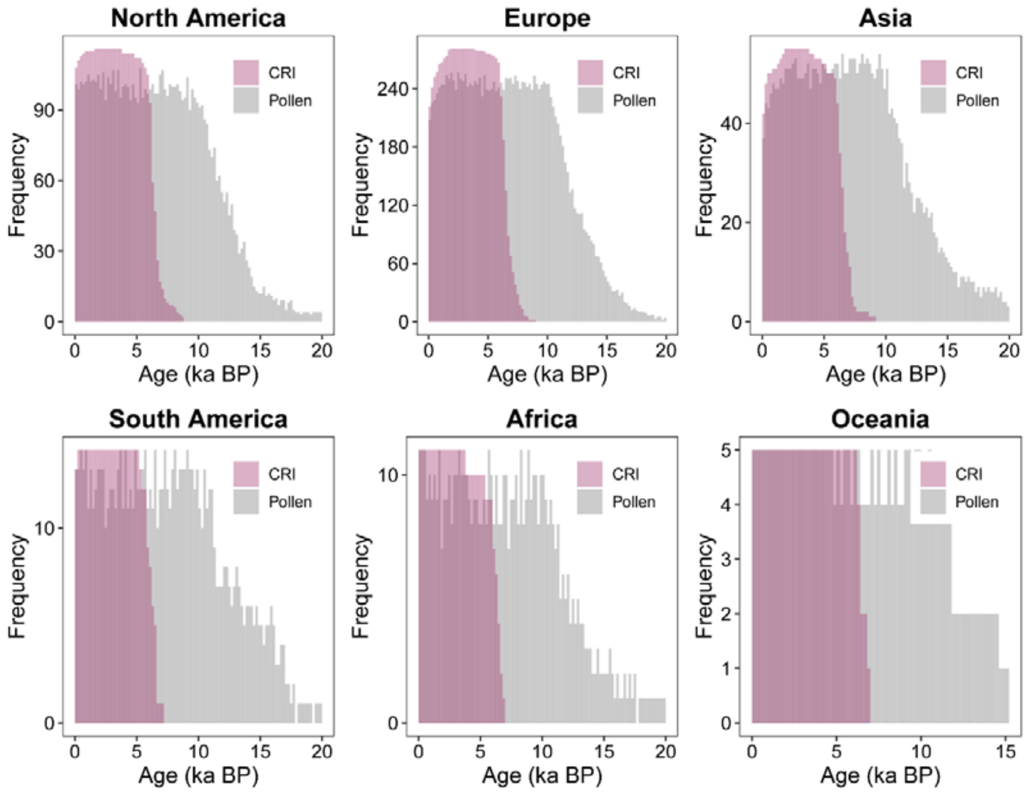


**Figure S1. Density of pollen and CRI sequences across continents.** Gray histograms indicate the number of pollen sequences per 200-year time interval over the past 20,000 years. Magenta-purple histograms depict the number of calculated CRI sequences per 200-year time interval during the past 8,000 years.

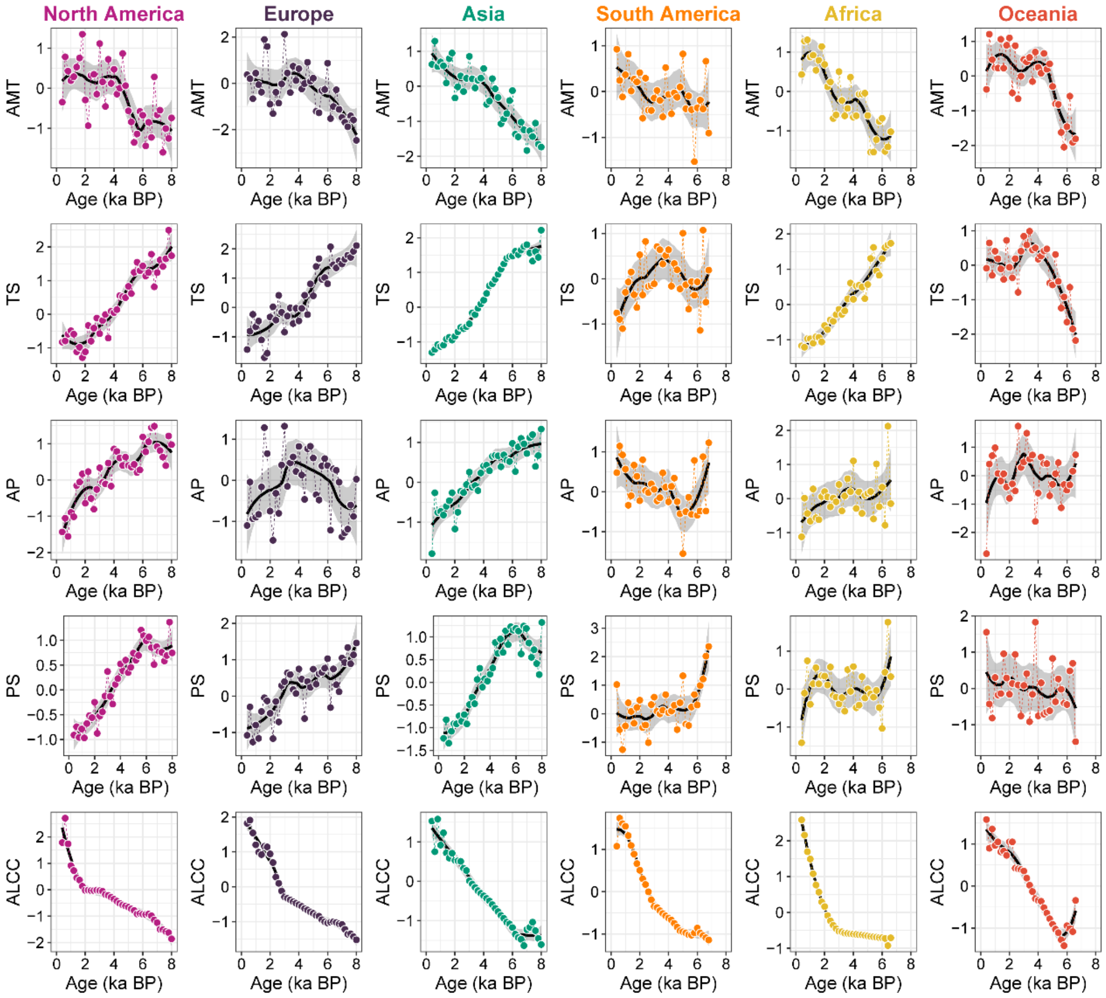


**Figure S2. Continental trends in climatic and land-use changes over the past 8,000 years.** Black solid lines represent (local polynomial regression) LOESS fits with a moderate smoothing window size (50% of data point used in each local regression). The shaded areas represent the 95% confidential intervals of the LOESS fits. The filled circles indicate the median values of each variable for every 200-year time interval. AMT, annual mean temperature; TS, temperature seasonality; AP, annual precipitation; PS, precipitation seasonality; ALCC, anthropogenic land-cover change.

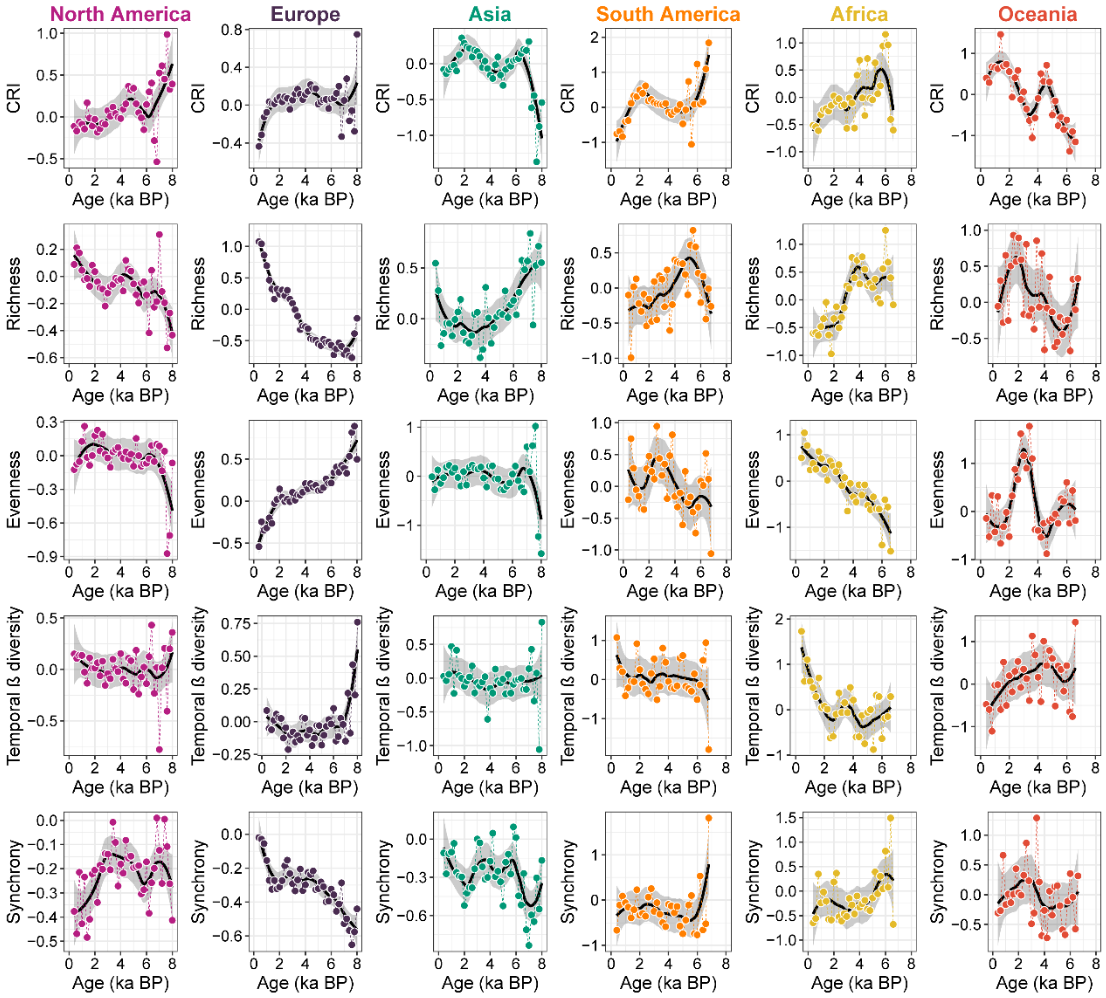


**Figure S3. Continental trends in CRI, taxonomic richness, evenness, temporal *β* diversity, and synchrony over the past 8,000 years.** Black solid lines represent (local polynomial regression) LOESS fits with a moderate smoothing window size (50% of data point used in each local regression). The shaded areas represent the 95% confidential intervals of the LOESS fits. The filled circles indicate the median values of each variable for every 200-year time interval.

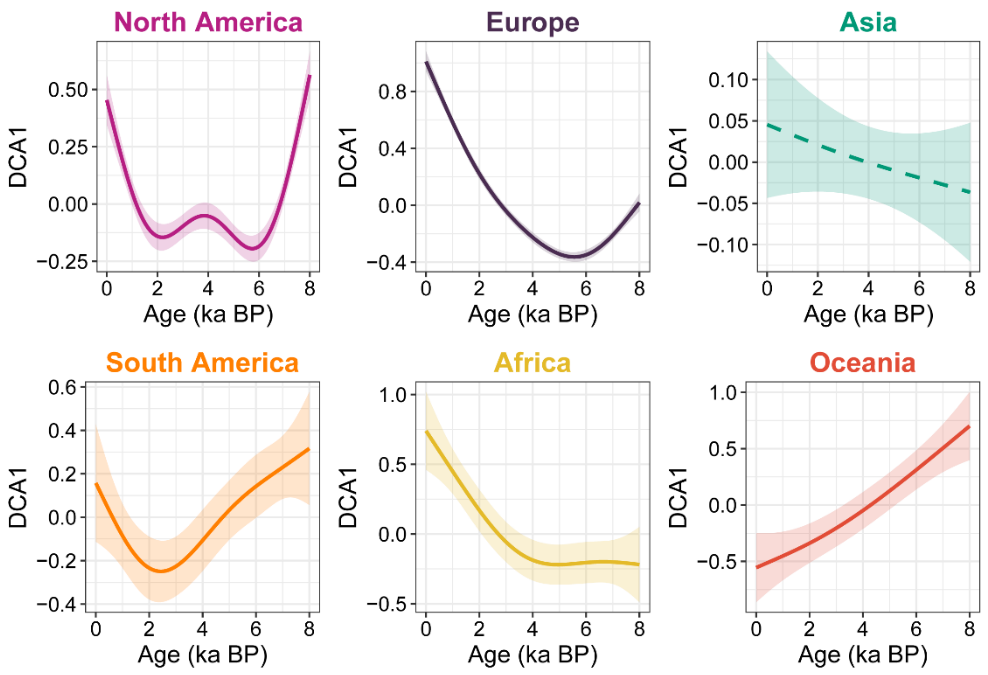


**Figure S4.** Continental trends of standardized first-axis scores of DCA (DCA1_st). Solid lines denote significant fits ($P < 0.05$) of generalized additive model (GAM), while dashed lines indicate non-significant relationship ($P > 0.05$). The shaded areas represent the 95% confidential intervals of the GAM fits.

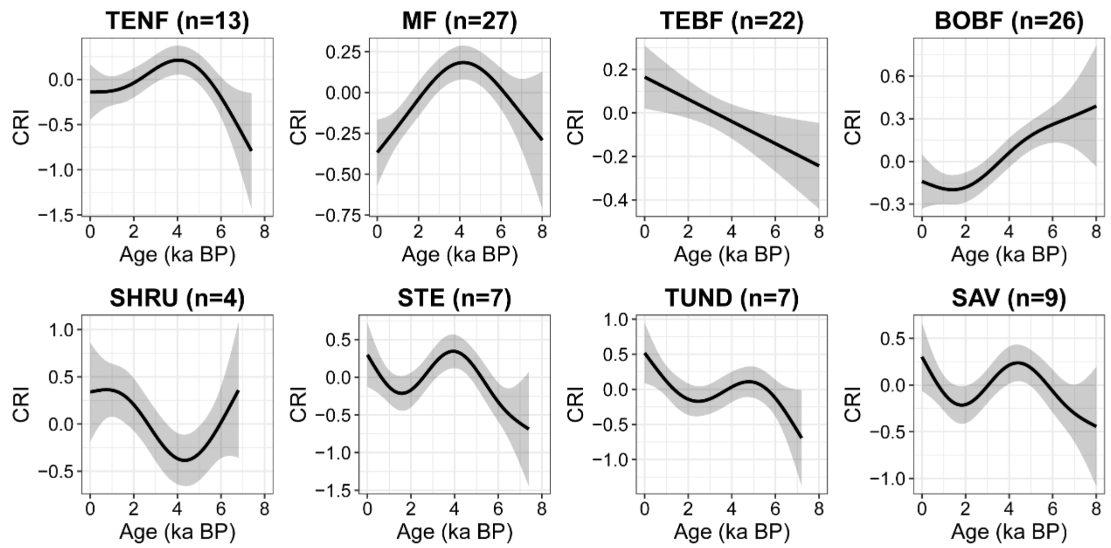


**Figure S5. CRI trends by biome across North America.** The shaded areas represent the 95% confidential intervals of the GAM fits. TEBF, temperate broadleaf forest/woodland; MF, mixed forest; BOBF, boreal deciduous forest/woodland; TENF, temperate needleleaf evergreen forest/woodland; STE, steppe/grassland; SHRU, shrubland; TUND, tundra; SAV, savanna; TRBF, tropical evergreen/deciduous forest/woodland; DESE, desert. Numbers in brackets denote the number of pollen sequences for modeling CRI trends.

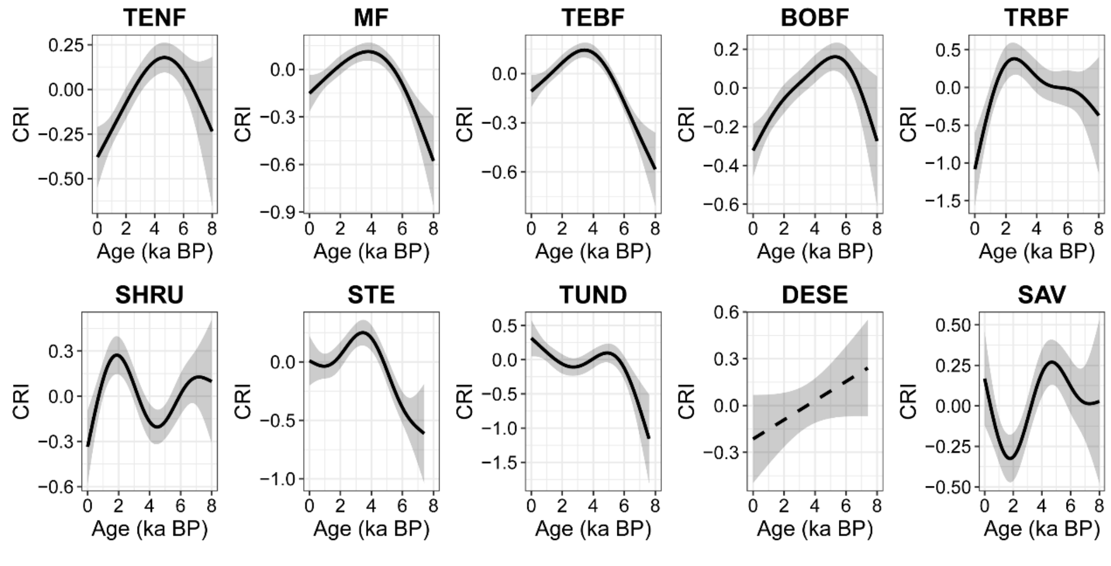


**Figure S6. Global CRI trends across different biomes.** All (generalized additive model) GAM fits demonstrate significant trends along the temporal gradient. Solid lines denote significant fits ($P < 0.05$) of generalized additive model (GAM), while dashed lines indicate non-significant relationship ($P > 0.05$). The shaded areas represent the 95% confidential intervals of the GAM fits. TEBF, temperate broadleaf evergreen forest/woodland; MF, mixed forest; BOBF, boreal deciduous forest/woodland; TENF, temperate needleleaf evergreen forest/woodland; STE, steppe/grassland; SHRU, dense shrubland; TUND, tundra; SAV, savanna; TRBF, tropical evergreen/deciduous forest/woodland; DESE, desert
